# Cell-cycle-dependent repression of histone gene transcription by histone H4

**DOI:** 10.1101/2024.12.23.630206

**Authors:** Kami Ahmad, Matt Wooten, Brittany N Takushi, Velinda Vidaurre, Xin Chen, Steven Henikoff

## Abstract

In all eukaryotes DNA replication is coupled to histone synthesis to coordinate chromatin packaging of the genome. Canonical histone genes coalesce in the nucleus into the Histone Locus Body (HLB), where gene transcription and 3’ mRNA processing occurs. Both histone gene transcription and mRNA stability are reduced when DNA replication is inhibited, implying that the Histone Locus Body senses the rate of DNA synthesis. In *Drosophila melanogaster*, the S-phase-induced histone genes are tandemly repeated in an ∼100 copy array, whereas in humans, these histone genes are scattered. In both organisms these genes coalesce into Histone Locus Bodies. We used a transgenic histone gene reporter and RNAi in Drosophila to identify canonical H4 histone as a unique repressor of histone synthesis during the G2 phase in germline cells. Using cytology and CUT&Tag chromatin profiling, we find that histone H4 uniquely occupies histone gene promoters in both Drosophila and human cells. Our results suggest that repression of histone genes by soluble histone H4 is a conserved mechanism that coordinates DNA replication with histone synthesis in proliferating cells.

## Introduction

The genome of eukaryotic cells is packaged into nucleosomes, where DNA is wrapped around histone octamers. In animal cells, the genes encoding canonical histone proteins are highly distinctive: These multi-copy genes are abundantly transcribed by RNA Polymerase II (RNAPII) during S phase of the cell cycle, and are the only protein-coding genes that produce transcripts without introns or 3’ polyadenylation^1^. Histone genes nucleate a distinctive body within the nucleus termed the Histone Locus Body (HLB), where specific transcription factors and RNA processing proteins localize. The HLBs of Drosophila and mammals share fundamental molecular components, including the Cyclin E/ CDK2-activated transcription cofactor Mxc/NPAT, 3’ mRNA stem-loops, Stem Loop Binding Protein (SLBP) and the U7 snRNP 3’-end processing machinery. Despite these cytological and compositional similarities, the histone genes are radically different in gene organization: In *Drosophila melanogaster* the five canonical histone genes (H1, H2A, H2B, H3 and H4), are arranged in a unit tandemly repeated ∼100 times at one locus^2^, whereas in humans 72 genes are scattered with a major cluster on Chromosome 6 and two minor clusters on Chromosome 1. These human genes are non-repetitive and embedded in euchromatic regions of chromosomes, while the tandemly repeated Drosophila genes are subject to heterochromatic silencing^3^, and it has been unclear how many of these 200 gene repeat units in a diploid cell are transcribed. By profiling both *D. melanogaster* and human histone genes, we aim to understand ancient conserved mechanisms of S-phase-induced histone gene regulation that have endured despite profound genomic and epigenomic changes.

Here, we have taken advantage of Drosophila histone gene rescue^4,5^ and fluorescently marked histone transgene reporters^6^ to test for histone gene derepression after knockdown of candidate repressors in the synchronized G2 phase gonial cells of testes. We discovered that reduction of histone H4 strongly derepressed histone gene expression outside of S phase, but no other candidate regulator had an effect. Using imaging, we showed that histone H4 localizes to the HLB in Drosophila cells; using CUT&Tag chromatin profiling, we precisely localized histone H4 to histone gene promoters, coinciding with peaks of Mxc/NPAT, initiating RNAPII and of active chromatin. Turning to human K562 cells, we observed similar cytological localization of histone H4 to the HLBs and coincident genomic localization of histone H4, NPAT, and RNAPII at active histone genes. These results imply a direct mechanism whereby excess histone H4 in cells buffers histone gene transcription to coordinate chromatin packaging with DNA replication.

## Results

### Chromatin features at active and silenced histone genes in Drosophila

As the histone genes in the Histone Locus Body (HLB) are repetitive, and normal cells contain both active and silenced histone genes, mapping of chromatin features to a genome assembly cannot distinguish which features are associated with which expression state. To address this, we profiled chromatin features in two genotypes: wildtype, where some of the ∼200 histone genes must be active while others are silenced, and the “*12XWT*” line, where the histone locus has been deleted and a construct carrying 12 copies of the histone repeat unit (HRU) rescues the flies^7^. We expect all copies of the histone genes to be active in this second genotype. By comparing chromatin profiles between these two genotypes, we infer the chromatin features of active and of silenced histone genes.

We dissected wing imaginal discs from male larvae as a sample of proliferating cells, and subjected them to CUT&Tag profiling^8^, generating >1 million reads mapped to the dm6 genome assembly for each sample (**Supplementary Table 1**). We first used antibodies to the HLB-specific transcription cofactors Mxc^9^ and Mute^10^. As expected, these two factors are localized to only the histone locus in the genome (**Figure 1a**). Signal for both factors is broadly dispersed across the *His3-His4* and *His2A-His2B* gene pairs of the 5 kb HRU, with peaks at the divergent promoters of each pair, and no signal over the adjacent *His1* gene (**Figure 1b**). These results are consistent with previous chromatin mapping of Mxc in Drosophila embryos^11^. The divergent *His4*-*His3* promoter region nucleates HLB formation^12^, and these binding profiles support the idea that Mxc binds at these sites in the histone locus and nucleates HLB formation. Further, the lack of Mute or Mxc signal over the *His1* gene is consistent with the idea that the linker histone gene is controlled by the CRAMP/ CRAMP1 transcription factor^13^.

**Figure 1.**
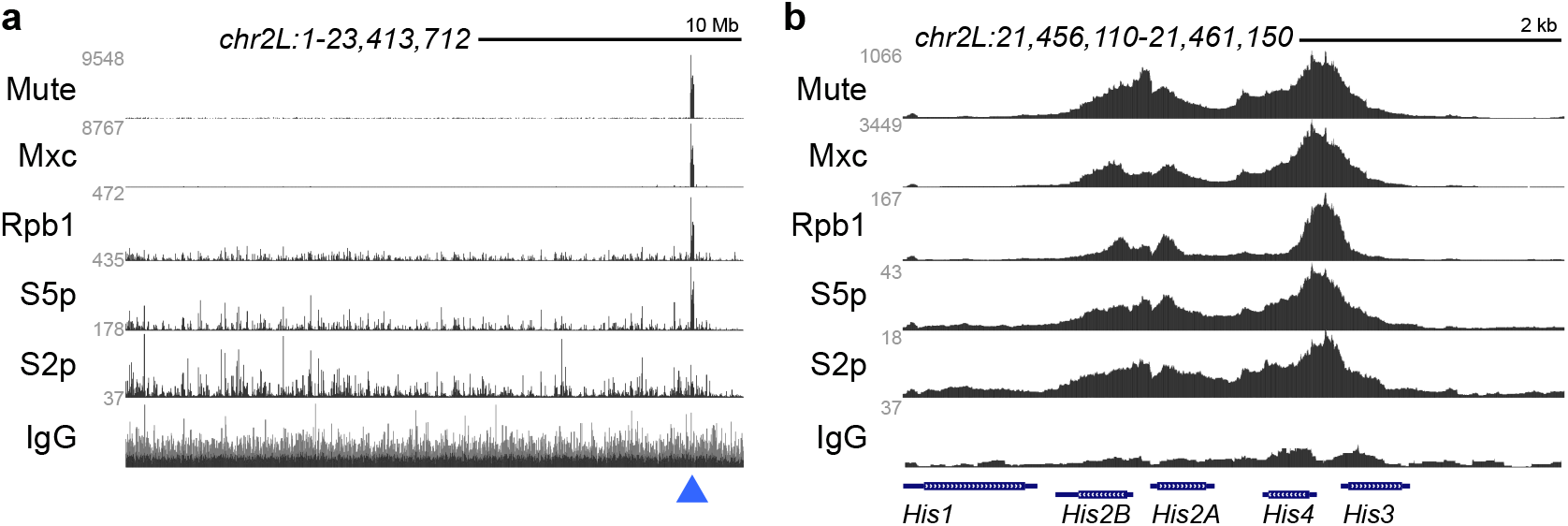
Chromatin factors and RNAPII at the histone locus in wing imaginal disc cells. **(a)** Browser tracks of chromatin factors and RNAPII isoforms across chromosome *2L* of Drosophila. The blue arrowhead marks the location of the histone locus. **(b)** Browser tracks of chromatin factors and RNAPII isoforms across one Histone Repeat Unit.

To assess transcriptional activity of the histone genes, we profiled multiple components and isoforms of RNA polymerase II (RNAPII) in larval wing disc samples. As expected, the RNAPII component unphosphorylated Rpb1, and phosphorylated initiating (RNAPII-S5p) and elongating (RNAPII-S2p) isoforms of Rpb1 mark the histone locus in wildtype cells (**Figure 1**). In fact, the histone locus is the major site of Rpb1 and RNAPII-S5p signal across the genome (**Figure 1a**), accounting for 1.8% and 2.5% of mapped reads, respectively, while only 0.2% of reads for the RNAPII-S2p isoform map to this locus. This is consistent with cytological description of enrichment for the unmodified and initiating forms of RNAPII at the HLB in other Drosophila cell types^14-16^. Note that the *His3-His4* and *His2A-His2B* have strong peaks of RNAPII-S5p and RNAPII-S2p isoforms at their promoters and across their gene bodies, indicating high transcription of these genes. In contrast, *His1* has only a low broad distribution of these polymerase isoforms across its length (**Figure 1b**), implying that it is expressed at lower levels.

We then compared signal counts for Mxc, Mute, and RNAPII between wing disc samples from wildtype and from *12XWT* flies. We calculated per-gene counts to adjust for the different numbers of copies in the two strains (100 copies/ genome in wildtype, 12 copies/genome in 12XWT). The per-copy sum counts of Mute and Mxc signal are higher in wildtype than in 12X flies, and proportional to the numbers of histone genes in these genotypes (**Figure 2a**). In contrast, signals for the RNAPII-S5p isoform are dramatically increased, and signal for the RNAPII-S2p isoform shows a slight gain. Thus, each histone gene in the 12XWT line must carry more RNAPII than in wildtype. There is no significant change in genome-wide gene expression (Supplementary Figure 1; Supplementary Table 3), consistent with the observation that the 12XWT rescue construct is sufficient to support viability and fertility^7^. The effect on chromatin features at histone genes implies that in wildtype some histone genes are active and others silenced, or alternatively that each gene is active an intermediate level. In either case, all histone genes in the 12XWT strain appear to be more heavily transcribed, presumably to support cell proliferation.

**Figure 2.**
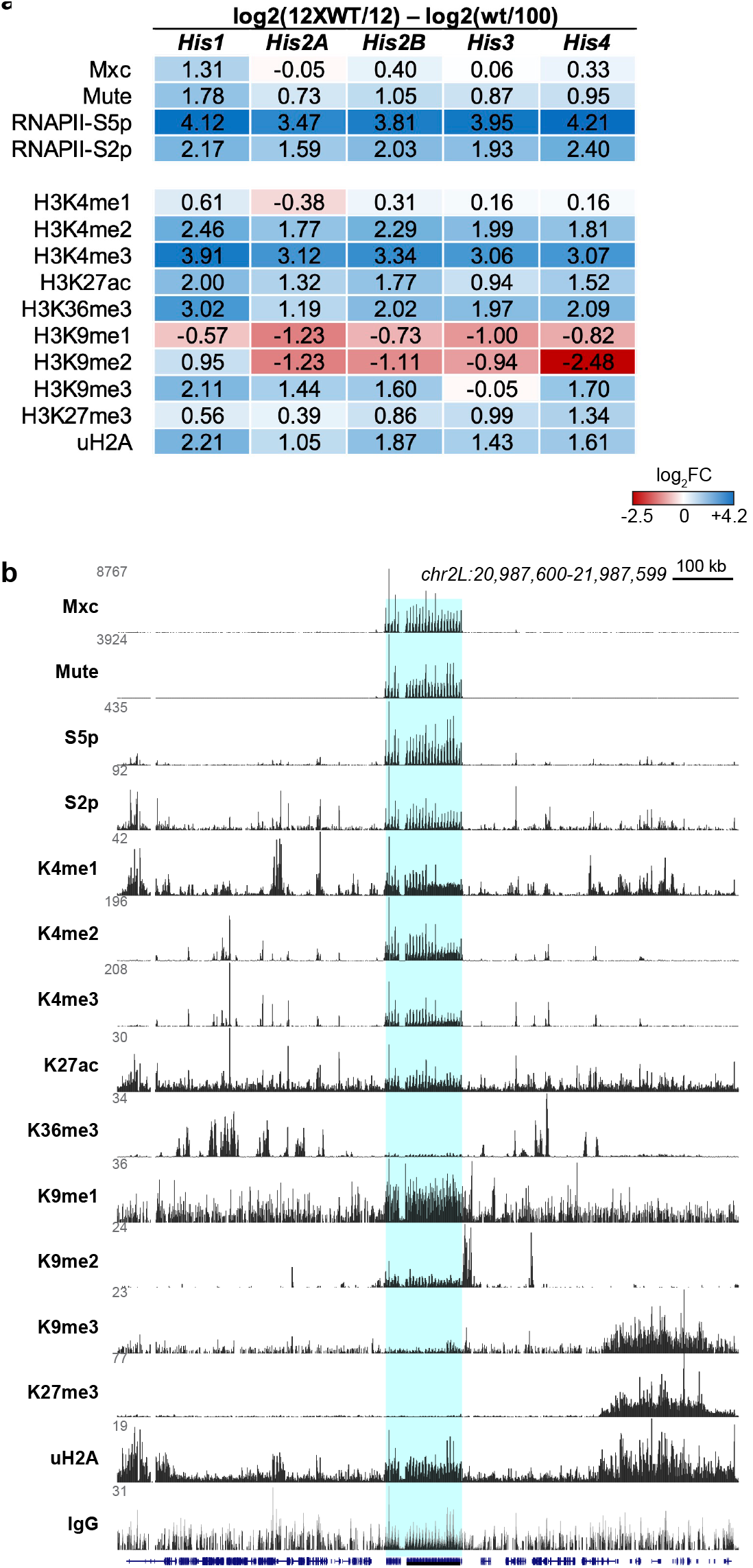
Histone modifications at the histone locus in wing imaginal disc cells. **(a)** Per-gene fold-change in chromatin features and histone modifications between the *12XWT* strain (12 histone gene copies/ genome) and wildtype (100 copies/genome). The H3K36me3, H3K9me3, and H3K27me3 modifications are absent from the wildtype and *12XWT* histone genes, and these ratios are due to minor changes in signal. **(b)** Browser tracks of histone modifications around the histone locus in wildtype wing imaginal disc cells. Blue shading marks the histone locus.

### Heterochromatic histone marks at silenced HLB genes

To determine histone modifications associated with histone genes, we first profiled five modifications associated with active gene expression^17^ (**Figure 2b**). In wildtype samples, all three methylation states of the histone H3K4 residue (H3K4me1, H3K4me2, and H3K4me3) and acetylation at histone H3-K27 (H3K27ac) are enriched at the wildtype histone locus at levels comparable to surrounding active enhancers and genes. In contrast, there is little detectable trimethylation of the K36 residue of histone H3 (H3K36me3), consistent with the intronless structure of histone genes^18^. Adjusting for copy number, there is a substantial per-gene copy gain only in the H3K4me3 mark in the 12XWT line (**Figure 2a**) and more moderate gains in the H3K4me1 and H3K4me2 marks. These changes are consistent with the higher average transcription of histone genes in this genotype.

We then examined five histone modifications typically associated with silencing across histone genes between the two Drosophila lines. Three modifications are strongly enriched at the histone genes in wildtype samples: mono- and di-methylation of histone H3 at lysine-9 (H3K9me1 and H3K9me2^19^), and ubiquitinylation of histone H2A at lysine-118 (uH2A^20^) (**Figure 2b**). The presence of these first two marks implies that histone genes have a partially heterochromatic character, likely due to the repeated genes, although the locus lacks tri-methylation of histone H3 at lysine-9 (H3K9me3^19^ (**Figure 2b**). The presence of uH2A at histone genes suggests that these are sites of PRC1 activity, although the canonical trimethylation mark of histone H3 at lysine-27 (H3K27me3^21^) mark of Polycomb-silenced domains is absent.

Since any histone modification associated with histone gene silencing should be present in wildtype but absent in the 12XWT line, we calculated per-gene copy changes for the three modifications that are above background levels across the histone locus (**Figure 2a**). Previous results have implicated H3K9 methylation in histone gene silencing^22,23^. Indeed, the per-gene copy coverage of the H3K9me1 and H3K9me2 marks drop in the 12XWT strain compared to wildtype. In contrast, per-gene density of the uH2A modification is slightly increased. These results support the idea that inactive histone genes are marked with mono- and di-methylation of the H3K9 residue, where the wildtype histone locus is a mixture of transcriptionally active histone genes mixed with silenced histone genes. This may be analogous to the functional organization of ribosomal RNA genes, where actively transcribed units are intermixed with silenced units^24^. Alternatively, individual histone genes may carry both active and repressive modifications that quantitatively adjust gene expression. In either case, a a partially active set of histone genes may allow cells to fine-tune histone production to the needs of cell proliferation and growth, the rates of which vary between tissues and life stages.

### A visual reporter for histone gene silencing

Extra histone repeat unit transgenes are repressed in proportion to the number of total histone genes in a genotype^25^ an effect that appears similar to the reduced expression of histone genes in wildtype. To visualize histone gene expression in living animals, we used HRU reporter constructs where either the *His3* or the *His2A* gene is fused to the octocoral *Dendra2* fluorescent protein coding sequence^6^. These constructs express fluorescently tagged histones in eggs and developing embryos^6^, and at low levels in proliferating cells of later stages such as in larval imaginal wing discs (**Figure 3a**). We reasoned that if these transgenes are partially repressed, then genetically interfering with histone gene silencing would produce more fluorescent protein. Indeed, the expression of the *His2ADendra2* HRU transgene is dramatically increased ∼37X in the *12XWT* background (12 HRU copies/genome) compared to its expression in wildtype in wing imaginal discs (**Figure 3b**). This implies that cells with reduced histone gene numbers sense a dearth of histones and specifically upregulate the histone locus to compensate and provide for chromatin duplication.

**Figure 3.**
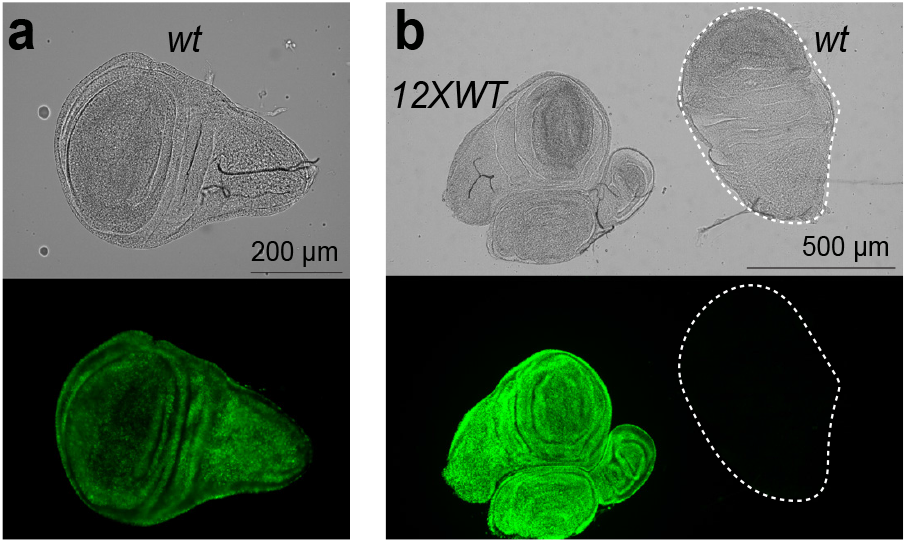
Expression of a *His2ADendra2* reporter Histone Repeat Unit in wildtype and *12XWT* strains. **(a)** Fluorescence of the *His2ADendra2* reporter in a wildtype larval wing imaginal disc, adjusted to display the weak fluorescence expressed from the reporter in this genotype. **(b)** Fluorescence of *His2ADendra2* in larval wing imaginal discs from wildtype (wt) with 100 HRU copies/genome and from *12XWT* larvae, with 12 HRU copies/ genome. Total fluorescence of *12XWT* wing imaginal discs is 37X that in the wildtype background (average summed fluorescence of wildtype discs 5,908,005 ± 1,096,058 a.u. *versus* 158,704 + 26,029 a.u. in the *12XWT* background).

We wished to identify the mechanism by which histone genes are repressed, but genetic reduction of such a factor might inhibit viability in the growing wing imaginal disc. Therefore, we turned to a non-essential tissue where we could still assess reporter expression with RNAi. The adult testis is dispensable for organismal viability. It also spatially organized so that the developmental and cell cycle stage of cells can be identified by their position in the tissue^26^. Germline stem cells are located at the apical testis tip and undergo four rounds of mitotic division in this proliferating zone, producing G2 phase spermatogonial cells. These then grow for ∼4 days before meiosis and sperm differentiation. We imaged these stages by dissecting and fixing testes from males carrying a *bamGAL4* driver with an inducible *UAS-RFP* transgene. This combination produces RFP specifically in gonial cells (**Figure 4a**). We immunostained these testes with antibodies to Mxc, a constitutive component of the HLB, and to phospho-Mxc/MPM2, which is catalyzed by Cyclin E/ Cdk2 kinase in S phase of the cell cycle^27^. The proliferating zone at the apical tip of the testis is marked with large HLBs containing phosphorylated Mxc and abut RFP-stained G2 phase cells (**Figure 4a**). Slightly more distal in the testis the Mxc-labeled HLB is divided into 2-4 smaller dots, consistent with the unpairing of homologous loci in this developmental stage^28^, and Mxc staining disappears from nuclei in later primary spermatocytes before meiosis.

**Figure 4.**
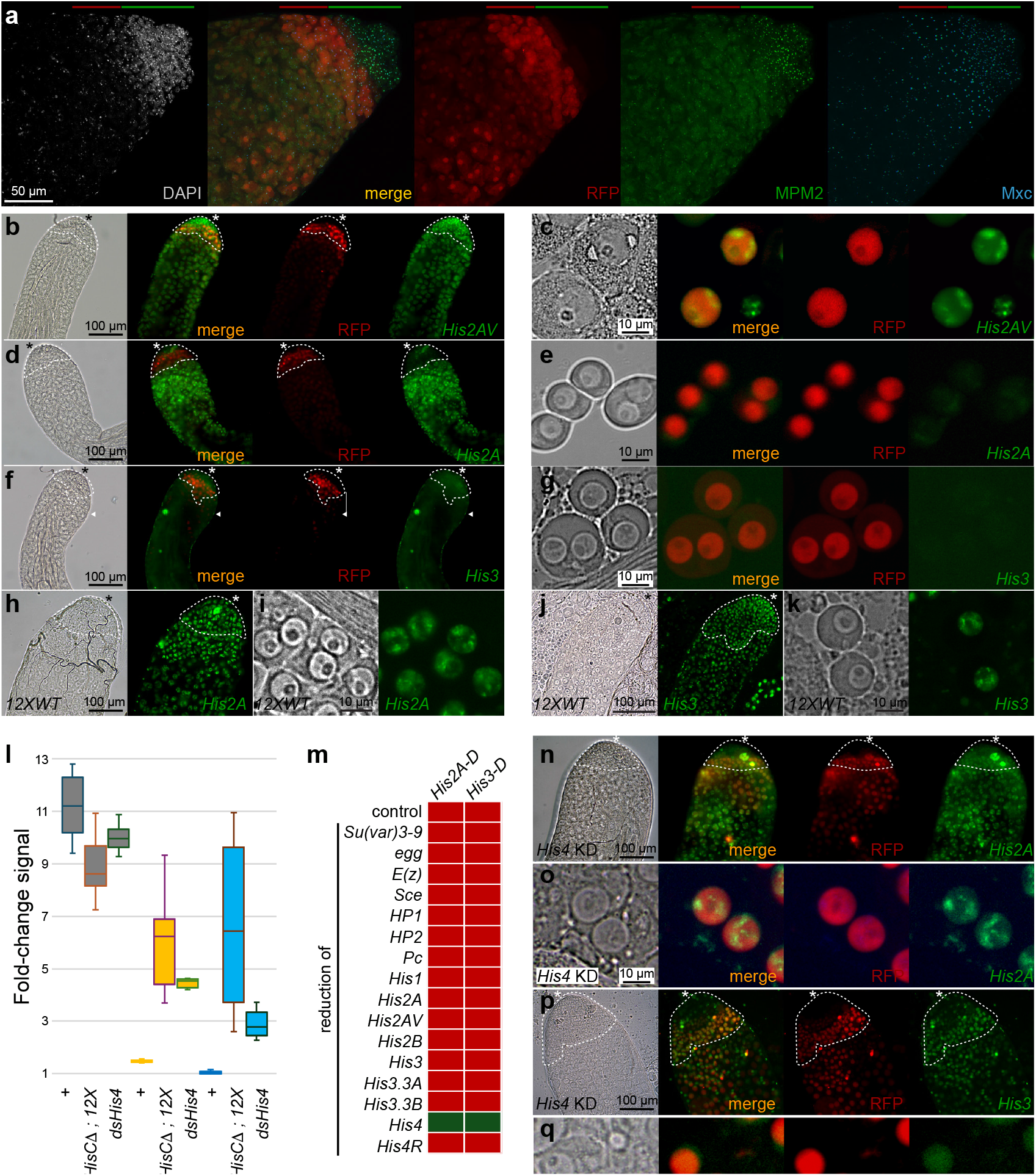
Silencing and derepression of extra histone genes in the Drosophila male germline. **(a)** The HLB in testis cells is marked by Mxc staining (blue). Cells in the proliferating zone at the apical tip of the testis are marked with phospho-Mxc detected by the MPM2 antibody at the HLB (green). Post-mitotic G2 phase germline cells are marked by *bamGAL4*-induced *UASRFP* expression (red), while the decondensed nuclei of later stages have low DAPI staining (grey). The proliferating zone of the testes is marked by the green bar, and the RFP-marked G2 gonial cells by the red bar. ((**b**,**c**) *His2AVDendra2* expression in wildtype testes. The apical tip is marked with an asterisk, the proliferating zone is outlined with white dashed lines on phase contrast images, and G2 phase gonial cells are identified with RFP. H2AVDendra2 fluorescence is apparent throughout the testis, including in the apical tip and proliferative zone, and gonial cells circled with a dotted line. H2AVDendra2 fluoresence is intense in the nuclei of isolated and squashed gonial cells (**c**), marked with *bamGAL4*-induced RFP expression. **(d**,**e)** *His2ADendra2* Histone Repeat Unit expression in wildtype males. Fluorescence is absent from the proliferating zone and from RFP-positive gonial cells but is present in later germline cells. **(f**,**g)** *His3Dendra2* Histone Repeat Unit expression in wildtype males. Staining is absent throughout the germline; the few fluorescent nuclei are in somatic cells of the testis sheath (arrowhead in (**f**). **(h**,**i)** Fluorescence of the *His2ADendra2* HRU reporter in 12XWT males. This line does not carry *bamGAL4* and *UAS-RFP* constructs; the proliferating zone and gonial cells were identified by position in the testis (dashedline), and in squashes gonial cells were identified by nuclear size and morphology. Strong nuclear H2ADendra2 fluorescence is apparent throughout the apical tip of the testis, in gonial cells, and in later stages. **(j**,**k)** Fluorescence of the *His3Dendra2* HRU reporter in *12XWT* males. This line does not carry *bamGAL4* nor *UAS-RFP*, and the proliferating zone and gonial cells were identified by position in the testis (dashed line), and in squashes gonial cells were identified by nuclear size and morphology. Strong nuclear H3Dendra2 fluorescence is apparent throughout the apical tip of the testis and in gonial cells. **(l)** Fluorescent intensities of gonial nuclei for the indicated genotypes with His2AVDendra2 (*H2AV-D*, grey), His2ADendra2 (*H2A-D*, yellow), or His3Dendra2 (*H3-D*, blue) reporters. **(m)** Results of tests for derepression of *His2ADendra2* and *His3Dendra2* Histone Repeat Unit reporters in testes with reductions in chromatin factors. *Su(var)3-9* was tested in *Su(var)3-9*^*1*^*/Su(var)3-9*^*2*^ homozygotes, all other factors were tested by *bamGAL4*-induced knockdown in testes. Red squares represent repression similar to wildtype, green squares indicate derepression. **(n-q)** Derepression of *His2ADendra2* (n,o, green) and of *His3Dendra2* (p,q, green) in cells with *bamGAL4*-induced RFP expression (red) and *His4* knockdown (KD). Fluorescence is absent in the apical tip of the testis but appears in the post-mitotic stage where knockdown occurs.

We first imaged fluorescence from a control *His2AVDendra2* histone variant gene (located on chromosome *3* outside of the histone locus), and this variant protein is abundant throughout the apical tip of the testis (**Figure 4b,c**). In contrast, the HRU transgenes are repressed in the testis. Little or no fluorescence from the *His2ADendra2* transgene is apparent in the very apical tip including in bam-positive gonial cells, but then weakly appears in later stages **(Figure 4d,e**). Fluorescent protein from the *His3Dendra2* HRU is undetectable throughout the testis, with signal only in the nuclei of somatic sheath cells (**Figure 4f,g**). We attribute the difference in pattern of expression between *His2ADendra2* and *His3Dendra2* transgenes to the deposition of histone H2A in post-mitotic cells, while histone H3 does not^29,30^ This repression is sensitive to the demand for histones, because animals with reduced numbers of histone genes and the *His2ADendra2* (**Figure 4h,i**) or *His3Dendra2* (**Figure 4j,k**) reporter HRUs now show intense fluorescence throughout the apical tip of the testis, including in gonial cells. Reduction of histone genes does not have a substantial effect on expression of the *His2AVDendra2* reporter (**Figure 4l**), indicating that this effect is specific to S-phase-induced histone genes. Thus, we infer that germline cells – like somatic cells – upregulate histone gene expression when these genes are limiting.

### Reduced histone H4 activates histone gene reporters

Previous work has implicated the H3K9-methyltransferase *Suppressor of variegation 3-9* (*Su(var)3-9*) in histone gene silencing^22,23^, and indeed we find that the histone locus is enriched for H3K9 methylation (**Figure 2b**). We constructed viable null *Su(var)3-9* flies from *trans*-heterozygous point mutant alleles^31^, however neither *His2ADendra2* nor *His-3Dendra2* reporters were derepressed in this background (**Figure 4m**). To identify mechanisms responsible for HRU repression, we targeted a selection of chromatin proteins for RNAi knockdown in the gonial cell stage. None of these knockdowns derepressed *His2ADendra2* or *His3Dendra2* reporters, including the histone H3 lysine-9 methyltransferase *eggless* (*egg* ^32^), the H3K9me2/3-binding proteins HP1 ^33^ or HP2 ^34^, the histone H2A ubiquitin ligase *Sex combs extra* (*Sce* ^35^), the PRC1 component *Polycomb* (*Pc* ^36^), or the histone H3 lysine-27 methyltransferase *Enhancer of zeste* (*E(z)* ^37^). We did not assess the effectiveness of these knockdowns in the testis so we cannot rule out that more complete elimination of these histone modifications might affect expression of histone gene reporters.

Since changes in histone gene number alter reporter expression, we used RNAi to knock down histones in gonial cells of the male germline. We tested knockdown of the linker histone gene *His1*, the core histone genes *His2A, His2B, His3*, and *His4*, of the histone variant genes *His2AV, His3.3A*, and *His3.3B*, as well as the orphan gene *His4R*. We tested histone knockdown constructs with two available histone GFP reporters and confirmed that these are effective in the testis (**Supplementary Figure 2**), but note that these males contain sperm and are fertile, implying that these knockdowns only partially reduce histone gene expression. The HRU reporters remained repressed in 8/9 of these histone knockdowns, but knockdown of the *His4* gene resulted in a ∼5-fold increase in *His2ADendra2* expression and a ∼3-fold increase in *His3Dendra2* expression in gonial cells (**Figure 4m-q**). Expression from a variant histone *His2AVDendra2* line is not affected by *His4* knockdown (**Figure l)**. These results imply that cells measure the demand for S-phase-induced histones based only on the H4 histone.

### Histone H4 localizes to the HLB in Drosophila cells

It is surprising that knockdown of only one histone modulates HRU silencing, since histones associate in dimers and in octamers as nucleosomes are assembled. However, some examples have been identified where monomeric histone exist^38^, and where a singular histone is used to measure chromatin in both Drosophila and in human cells^39,40^. To test if histone H4 localizes to the HLB on its own, we examined the localization of different histones within the male germline using inducible GFP-tagged constructs^41^. Induction of a tagged H3 or tagged H3.3 histones in gonial cells broadly labels the nuclei of these cells (**Figure 5a,b**). In contrast, tagged H4 histone shows a distinct subnuclear pattern, with one major dot in each nucleus (**Figure 5c**). This dot coincides with Mxc protein at the HLB in gonial cells (**Figure 5d**). Since the *bamGAL4*-induced H4GFP construct is not expressed in proliferative zone cells, we stained testes with an antibody to histone H4. This revealed an H4 dot in proliferating zone nuclei, including most cells actively undergoing DNA replication (**Figure 5e**) and interphase cells **(Figure 5f)**. Only 7% (1/14) of prophase cells show the dot (**Figure 5g**), consistent with the partial disassembly of the HLB in mitosis^9^. Although histone H4 is incorporated throughout chromatin, the lack of widespread staining suggests that this antibody recognizes a part of the histone that is buried within nucleosomes, and that a soluble, non-nucleosomal histone protein is in the right place to directly affect histone gene expression both in S phase and in gap phase cells.

**Figure 5.**
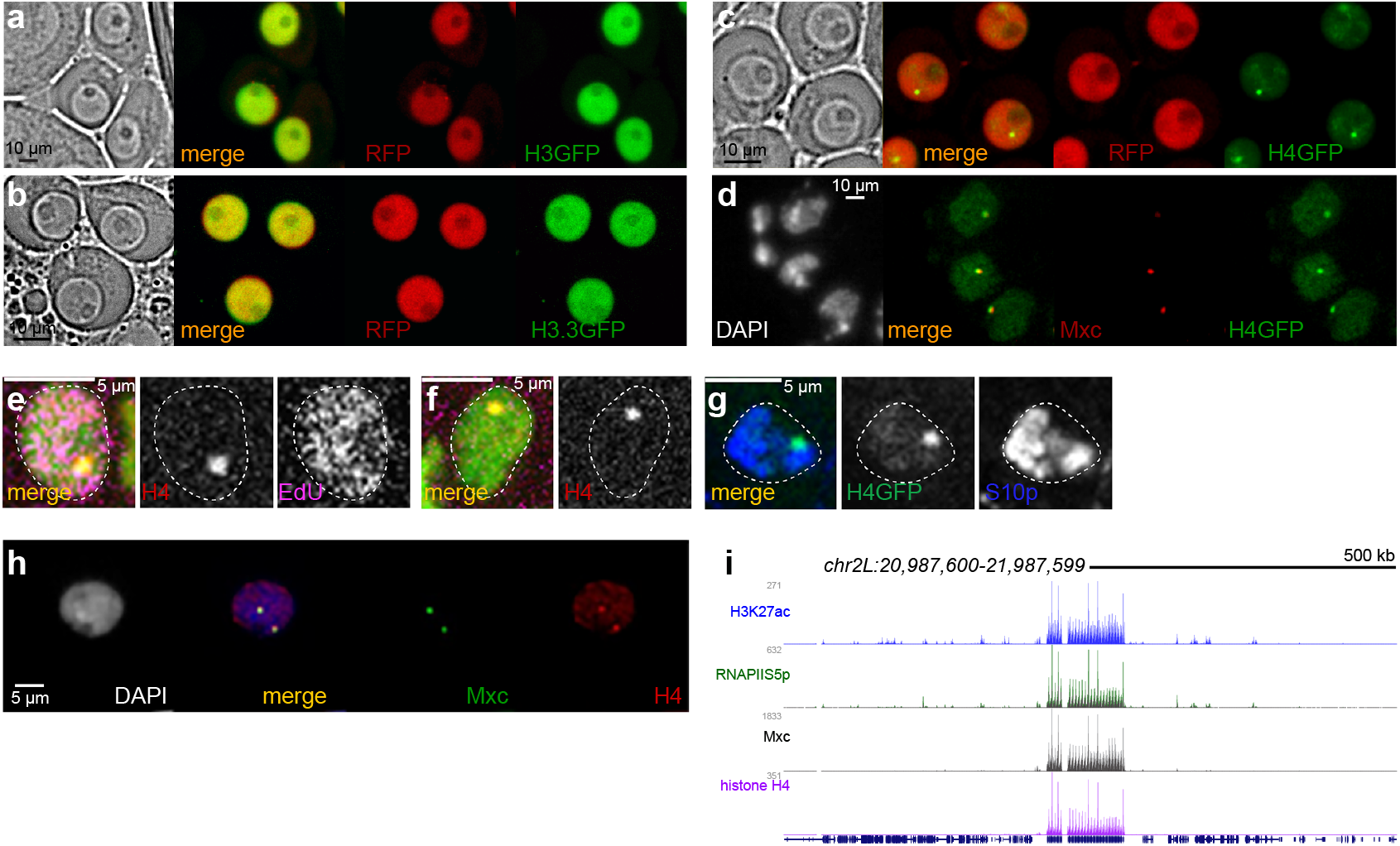
Histone H4 is a component of the Histone Locus Body in male germline cells. (**a-c**) Fresh squashes of testes with *bamGAL4*-induced expression of GFP-tagged histones (green) Induced RFP expression (red) marks G2 phase gonial cells. Histone H3-GFP **(a)** and histone H3.3-GFP **(b)** broadly label the nuclei of gonial cells. In contrast, induced histone H4-GFP **(c)** labels one bright dot with a low broad background in G2 phase nuclei. **(d)** The induced histone H4-GFP (green) dot coincides with the HLB, marked by Mxc staining (red). **(e**,**f)** Representative spermatogonial cells labeled with EdU (pink) to mark nuclei with ongoing DNA replication or in gap phase. DAPI staining is in green. 94% (47/50) S phase cells show focal staining of histone H4 (red) and this dot is also visible in gap phase cells. **(g)** Representative spermatogonial cell marked with the M phase epitope H3S10p (blue). The histone H4GFP (green) dot is visible in only 7% (1/14) of mitotic cells. **(h)** *Kc167* nucleus stained with antibodies to histone H4 (red) and Mxc (green). Histone H4 is enriched in HLBs. **(i)** Browser tracks of histone modifications around the histone locus in *Kc167* cells. The histone locus is enriched for the active H3K27 acetylation modification (blue), the RNAPII-S5p isoform (green), Mxc (black), and for histone H4 (purple).

We examined Drosophila cultured *Kc167* cells to determine if histone H4 localizes to the HLB in other cell types. In immunostained samples, histone H4 staining colocalizes with Mxc at HLBs (**Figure 5h**). In chromatin profiling, histone H4 shows high signal only at the histone locus, coincident with Mxc, H3K27 acetylation, and RNAPII-S5p signal (**Figure 5i**). As *Kc167* cells are derived from somatic embryonic cells, this implies that non-nucleosomal histone H4 is a general component of the HLB.

### Loss of histone H4 alters HLB activity

The HLB is enriched for the initiating RNAPII-S5p isoform in embryos and in the female germline^15,16,42^. In the testis, RNAPII-S5p is distributed through nuclei and forms a bright focal spot at the HLB in proliferating zone nuclei with bright phospho-Mxc staining, and weaker focal staining in RFP-positive G2 phase gonial cells (**Figure 6a-c**). This is the major isoform of RNAPII engaged at histone genes, as staining for the RNAPII-S2p isoform is broadly distributed throughout nuclei with no greater enrichment at the HLB (**Figure 6d,e**). By the early primary spermatocyte stage the HLB is no longer enriched for any RNAPII isoform. We infer that active histone loci in the proliferating zone are engaged with high levels of the RNAPII-S5p isoform, and a reduced amount of this isoform persists in G2 gonial phase cells when histone genes are no longer expressed. Overall, the HLB in the testis progresses from having high levels of engaged RNAPII in the proliferating zone, to less engaged and non-transcribing RNAPII in G2 phase gonial cells, and to loss of RNAPII in primary spermatocytes. Dissolution of the HLB occurs in later stages, as Mxc staining is eventually lost, as cells do not need histone gene expression as they proceed to meiosis and sperm differentiation. This progression and the spatial arrangement of the testis is an easily tractable setting to follow changes in the HLB.

**Figure 6.**
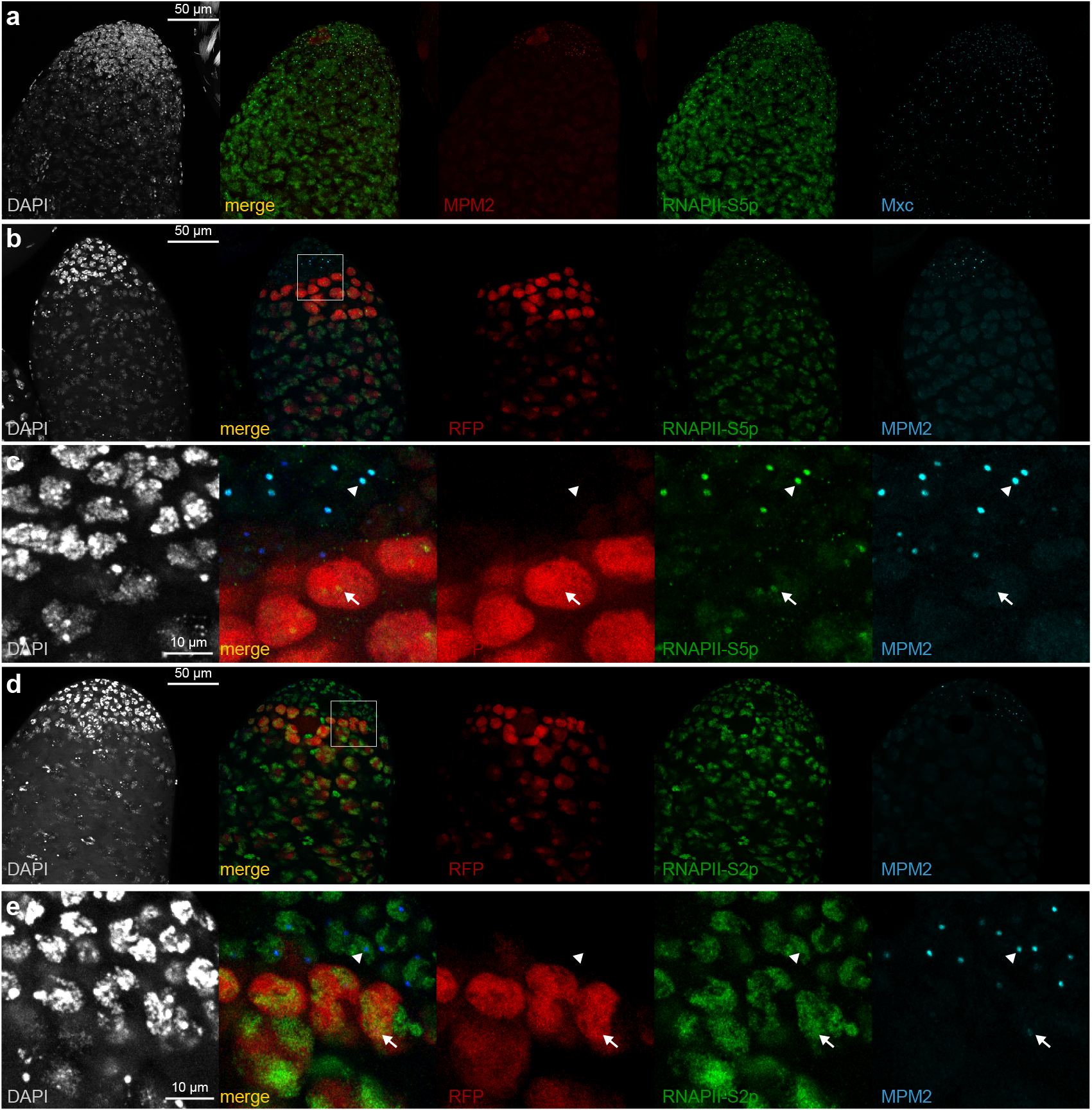
RNAPII isoforms at the HLB in the Drosophila testis. **(a)** RNAPII-S5p (green) stains the HLB in the proliferating zone marked by phospho-Mxc staining (red) and and the HLB in more distal cells. Mxc staining (blue) marks the HLB. **(b)** The proliferating zone is marked with phospho-Mxc (blue) and the G2 phase gonial cells by *bamGAL4*-induced RFP (red). RNAPII-S5p staining (green) strongly stains the HLB in the proliferating zone, and more weakly stains the HLB in G2 phase cells. **(c)** zoom of the boxed area in (b). The arrowhead points to an HLB in proliferating zone, and the arrow to an HLB in a G2 phase cell. **(d)** RNAPII-S2p staining (green) strongly stains nuclei in the proliferating zone and in RFP-labeled G2 phase cells, but is not enriched in HLBs. **(e)** zoom of the boxed area in (d). The arrowhead points to an HLB in proliferating zone, and the arrow to an HLB in a G2 phase cell.

The *bamGAL4*-induced H4-GFP dot is most intense in gonial cells with reduced MPM2 staining (**Figure 7a,b**). In wildtype testes, the most intense spots of RNAPII-S5p staining are in HLBs of proliferating cells with high MPM2 staining, while HLBs of gonial cells show a ∼60% reduction in RNAPII-S5p staining (**Figure 7c,e,g**). This suggests that histone H4 has a role in limiting histone gene expression. We then examined testes where *His4* was knocked down in gonial cells. Knockdown ablates H4 staining in the HLB (**Supplementary Figure 2**) and, in contrast to wildtype controls RNAPII-S5p staining increases by ∼30% (**Figure 7d,f,g**). These defects implicate histone H4 in the switch from active to inactive forms of the HLB.

**Figure 7.**
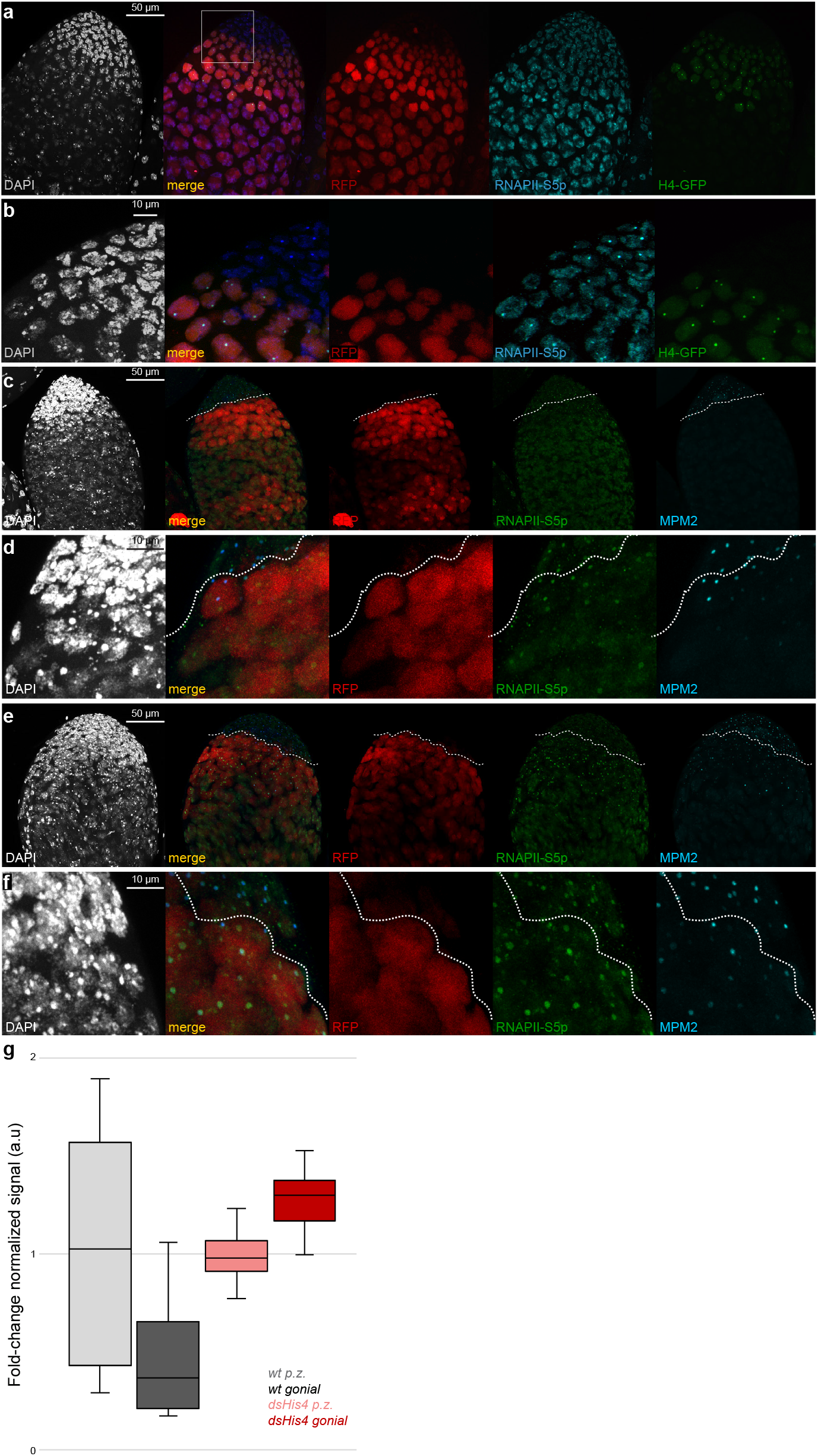
Histone H4 localizes with reduced RNAPII in HLBs. **(a)** Immunostaining for RNAPII-S5p (blue) in the apical tip of a testis with *bamGAL4*-induced expression of histone H4-GFP (green). G2 phase gonial cells are marked by induced RFP expression (red). The intense dot of H4-GFP coincides with focal RNAPII-S5p signal in G2 phase gonial cells. **(b)** zoom of the box marked in (a) showing co-staining of HLBs with RNAPII-S5p and H4-GFP in RFP-positive G2 phase gonial cells. RNAPII staining in these cells is lower than that in more apical cells in the proliferating zone. **(c-f)** Testes with *bamGAL4*-induced RFP expression (red) marking G2 gonial cells and stained for phospho-Mxc (blue) and the RNAPII-S5p isoform (green). Dashed lines demarcate the proliferative zone from G2 phase cells based on RFP expression. A wildtype testis **(c**,**d)** with high phospho-Mxc in the proliferative zone and low staining in G2 phase cells. RNAPII-S5p signal in the proliferative zone is high at some HLBs and moderate at others, and moderate or low at HLBs in the RFP-marked cells. **(e**,**f)** A testis with knockdown of *His4* (KD). High phospho-Mxc signal is apparent in the proliferative zone and occasionally persists in the RFP-labeled gonial cells. The RNAPII-S5p signal is present at HLBs of proliferative zone cells, and more intense at **g** _2_ the HLBs in gonial cells. **(g)** Quantitation of RNAPII-S5p signal in HLBs in proliferative zone (p.z.) and in gonial nuclei for the indicated genotypes. RNAPIIS5p staining in wildtype testes is high in proliferative zone cells and decreases in gonial cells. In testes with *His4* knockdown, RNAPII-S5p is high in the proliferative zone and increases in gonial cells. The intensity of RNAPII-S5p staining in knockdown gonial cells is ∼2X that of gonial cells in wildtype.

### Histone H4 localizes to active histone gene promoters in human cells

Since histone genes are repeated and we cannot distinguish between gene copies, we cannot define whether histone H4 binds all histone genes or only active or silent genes in Drosophila. But histone H4 is an ancient protein, and if it plays a role in limiting histone gene expression we expect HLB localization to be conserved across species. In the human genome the multiple copies of histone genes are not in a repeat array; instead they are comparatively separated and scattered across four clusters, termed *HIST1-4* ^43^, and these aggregate into HLBs containing the transcription cofactor NPAT, the human homolog of Mxc ^44,45^. Indeed, immunostaining of human K562 cells reveals that NPAT and histone H4 colocalize at HLBs (**Figure 8a**). We then profiled the distribution of histone H4 and the active histone modification H3K27 acetylation in K562 cells and compared these to previously published profiling of NPAT ^8^ and RNAPII-S5p ^46^. As expected, 64 canonical histone genes in the *HIST* on chromosome *6* are marked with both RNAPII and with H3K27mac modifications, identifying them as active genes (**Figure 8b-d**). Each of these active histone genes are also marked with NPAT and histone H4 (**Figure 8b**), and at high resolution these four chromatin features coincide at promoters (**Figure 8c**). In contrast, the 8 non-transcribed canonical histone genes lack both NPAT and histone H4 (**Figure 8b,d**), as do the 5 active histone variant genes that are not S phase-regulated (**Figure 8d**). Actively transcribed non-histone genes neighboring *HIST* clusters also lack NPAT and histone H4 signal (**Figure 8b**). These results implicate histone H4 in the S phase regulation of canonical histone gene promoters in human cells.

**Figure 8.**
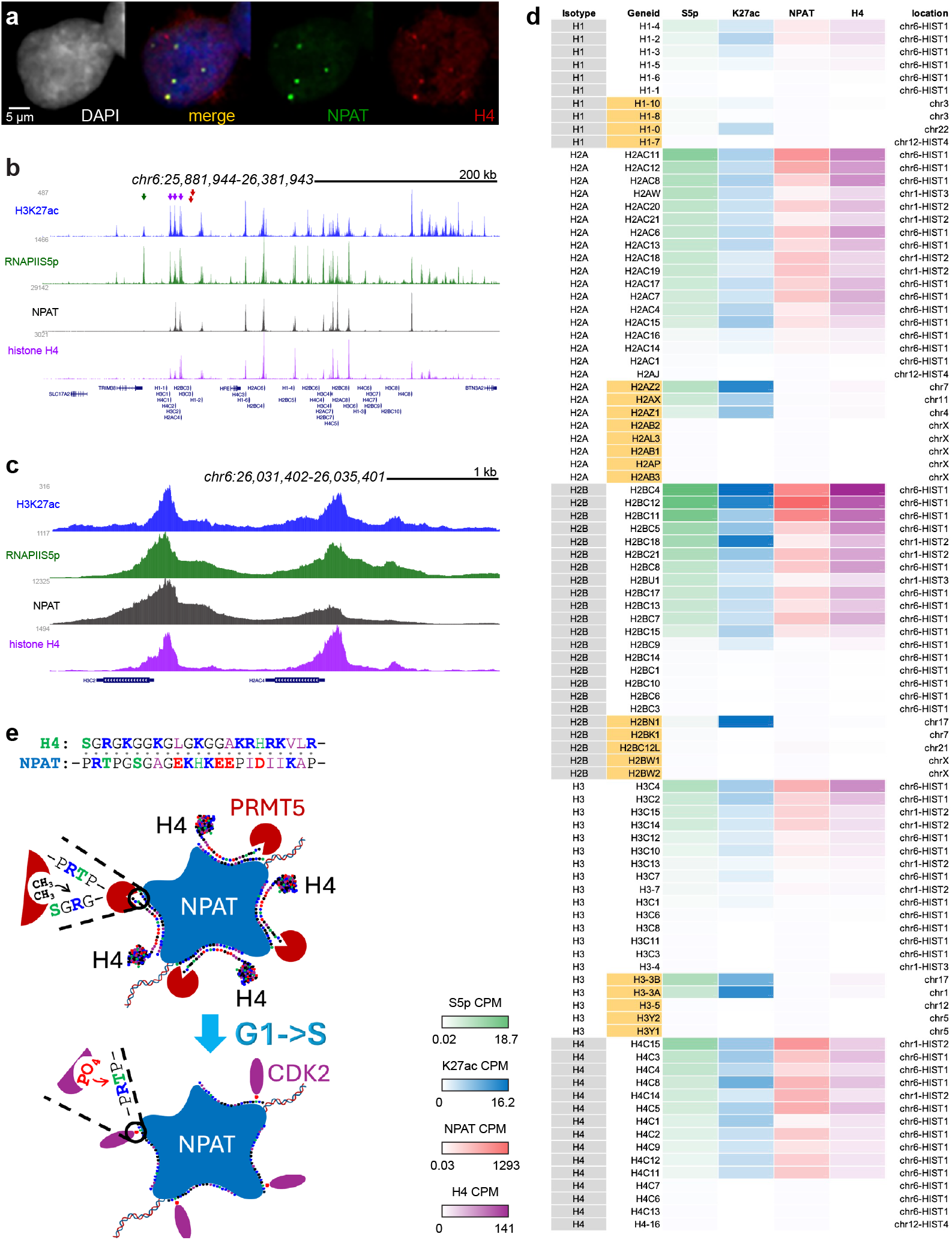
Histone H4 localizes to the promoters of active histone genes in human K562 cells. **(a)** Immunostaining human K562 cells for the HLB factor NPAT (green) and histone H4 (red). Anti-histone H4 signal localizes to each of the multiple HLBs in the nucleus. **(b)** Distribution of histone H3K27acetylation, RNAPII-Sp5, NPAT, and histone H4 across a portion of the *HIST1* histone cluster on chromosome *6*. Purple arrowheads mark three active canonical histone genes on one side of this cluster, the red arrowheads marks two inactive histone genes, and the green arrowhead marks the promoter of an adjacent active non-histone gene. NPAT and histone H4 coincide only at active histone genes, and are absent from inactive histone genes and from active non-histone genes. **(c)** Zoom showing coincidence of the H3K27 acetylation mark, RNAPII-S5p, NPAT, and histone H4 at the promoters of two active histone genes. **(d)** Summary of summed signal for chromatin features at 94 histone genes in the human genome. Variant histone genes are labeled in orange. Each histone isotype and variant is ordered by expression (RNAPIIS5p signal). **(e)** Model for S phase activation of histone gene expression. Top: Repression of NPAT by Histone H4 is mediated by weak van der Waal interactions between the basic N-terminal tail of histone H4 and unstructured regions of NPAT bound to histone gene promoters. Schematic shows that at the point of cell-cycle commitment, PRMT5 (brown) unpeels the H4 N-terminus and di-methylates H4R3, permanently releasing it from NPAT. This exposes the unstructured hydrophilic regions of NPAT, which is then activated by the CyclinE/CDK2 (purple) master regulator of entry to S-phase at Threonine-1270 and at least three other mapped positions.

## Discussion

The invention of eukaryotic chromatin required coordination of histone protein synthesis with DNA replication in S phase cells. DNA replication and histone synthesis must be coupled because even small imbalances are detrimental, given the large amounts of chromatin duplicated each S phase. Underproduction of histones will result in incomplete chromatin packaging, and this leads to exposure and damage of new DNA^47^ Histone overproduction results in chromosome loss^48^ and is cytotoxic^49^. Feedback between these two processes provides just enough histones to package newly replicated DNA^1,50^. Feedback control implies that S phase cells measure both ongoing DNA replication and the need for more histones. Histone production is modulated through two main controls: CDK2-catalyzed phosphorylation of Mxc/ NPAT induce transcription canonical histone genes when cells commit to S phase^51^, and cell cycle-regulated associations between histone mRNA processing factors stabilize transcripts in S phase^52^; both of these processes take place in HLBs^53^. S-phase-induced histone mRNAs are the only protein-coding transcripts that are not polyadenylated in animals; instead, these transcripts have a terminal stem-loop structure that is bound by stem-loop binding protein (SLBP), directing 3’ processing^52^. Mathematical modeling has suggested that feedback from soluble histone pools is necessary for precise coupling between DNA replication and histone synthesis^54^. Our observations suggest a simple model where soluble histone H4 protein directly represses histone gene transcription. Ongoing DNA replication and chromatin packaging use up soluble histones, but once DNA replication ceases, soluble histones including histone H4 accumulate. Monomeric histones have been observed in cells^38^, and so a monomeric histone H4 might directly interact with NPAT at histone gene promoters. The majority of the NPAT and Mxc proteins are composed of intrinsically-disordered domains (IDRs) interspersed with structured domains, and both are important for self-associations and function^55,56^. We propose an interaction between the strong positive charge of the unmodified H4 N-terminal tail (N-SGRGKGGKGLGK-GGAKRHRKVLR) and the strong negative charge of the unstructured regions of phosphorylated NPAT. Weak, transient charge interactions have been observed to drive chromatin binding of transcription factors^57^, and in the case of H4-NPAT would make H4R3 available for access by PRMT5 for arginine di-methylation and H4 release from NPAT.

The atypical 3’ ends of histone genes appears to be the ancestral organization of S-phase-induced histone genes throughout Eukaryota, because both stem-loop mRNA structures and SLBP homologs have been identified in protozoa at the base of the eukaryotic tree^58^. The evolutionary origin of this system was unclear, but recently similarities to 3’ processing of transcripts in bacteria have been pointed out^59^. Thus, the histone 3’ processing system appears to be one of the few relics in eukaryotic genomes of their origin and may explain why these genes are sequestered in their own nuclear body. The eukaryotic histones themselves were derived from bacterial or viral proteins in the last eukaryotic common ancestor^60^, with eukaryotic histone H4 being a sister lineage to both the HMfB archaeal histones and the doublet H4-H3 histones of the Nucleocytoviricota giant viruses^61^, which later diversified into the four core histone subtypes^62^. Thus, it is conceivable that histone H4 has been used as a negative regulator of core histone gene expression all this time. Indeed, histone H4 is distinctive in that variants for this isotype rarely occur across eukaryotic evolution. One exception is a variant histone H4 encoded by some symbiotic bracoviruses; braconid wasps harboring this virus inject viral DNA into host moth larvae, where production of the variant histone suppresses host histone H4 mRNA production^63^. This unusual variant appears to have weaponized the normal negative feedback loop of histone H4 on histone gene regulation for parasite life history.

Histone gene regulation has been implicated in cancer progression in human patients, and histone overproduction is predictive of cancer malignancy^64^. There are multiple theories to explain the initiation of a cancer, invoking genetic mutation, changes in epigenetic marks, and defects in developmental signaling. Although the relative importance of these theories is now debated^65,66^, in any scenario progenitor cells must maintain a relatively undifferentiated state and proliferate. We have documented that widespread chromosome arm aneuploidies are common in cancers and number of arm losses scales with malignancy^67^. Such aneuploidies in tumors were suggested to inhibit cellular differentiation and thereby trap anaplastic cells in a proliferative stage^68,69^. We have proposed that these events are causally linked: that histone overexpression during S phase compromises histone variant-based centromere assembly and also accelerates cell proliferation; these effects would directly cause mitotic chromosome errors and result in aneuploidies^67,69^. In support of this scenario, a recent study identified reduced cell cycle duration as the only common feature of multiple distinct cancers, showing that tumorigenesis could be blocked by mutations affecting CDK2 activity^70^. CDK2 is the kinase that phosphorylates Mxc/NPAT and histone gene activation, linking conserved HLB regulation described here to a deeper understanding of cancer. Further, a second recent study has demonstrated that inhibition of PRMT5, which symmetrically di-methylates histone H4 arginine-3, results in rapid hist gene repression^71^. While the effects of anti-cancer PRMT5 inhibitors have been attributed to mRNA splicing defects, the very rapid effect on histone gene expression implies that the interaction surface between histone H4 and NPAT is the primary target of these anti-cancer drugs (**Figure 8e**). As such, we anticipate that further insights into the direct interaction we propose will provide further opportunities for intervention.

## Methods

### Fly strains

All crosses were performed at 25^°^C. All mutations and chromosomal rearrangements used here are described in Fly-base (http://www.flybase.org). The *w*^*1118*^ strain was used as a wildtype control. The *12XWT* strain is *w*; Δ*His*^*C*^; *12XWT* ^5^. The HRU reporter *His3Dendra2* was previously described^6^, and the *His2ADendra2* reporter was constructed similarly. Inducible histone lines *UAS-H3-GFP*, and *UAS-H3.3-GFP* were previously described^41^, and the *UAS-H4-GFP* construct was injected into fly embryos for P element transformation^73^ by BestGene Inc. (Chino Hills CA). A similar UAS-H4-eGFP construct used here for some experiments was previously published^74^. Additional constructs used for cytological characterization were *y w P[bam-GAL4:VP16,w*^*+*^*]1/Y*; *P[UAS-RFP,w*^*+*^*]2*. Inducible knockdown constructs and Bloomington Drosophila Stock Center IDs for histones and chromatin regulators are listed in the *Key Resources Table*.

### Antibodies

Antibodies used for CUT&Tag profiling and for immunocytology are listed in the *Key Resources Table*.

### Imaging fresh tissues

Dissected tissues from larvae or adults were mounted in PBS on slide and imaged by epifluorescence on an EVOS FL Auto 2 inverted microscope (Thermo Fisher Scientific) with a 10X, 20X, or 40X objective. Pseudo-colored images were adjusted and composited in Adobe Photoshop and Adobe Illustrator. For measuring signal intensities of imaginal discs, we used Photoshop to select the entire disc by phase contrast imaging and summed the GFP fluorescence pixel intensity of that area in unsaturated images with identical camera settings for genotypes. For measuring signal intensities of nuclei, we used Photoshop to select gonial cell nuclei by RFP staining or by phase contrast nuclear morphology, summed the Dendra2 fluorescence pixel intensity of that area, and calculated the mean pixel intensity in unsaturated images. We normalized pixel intensities by dividing by the mean background signal in that image, so that no signal fluorescence is 1.

### Imaging immunostained testes

Testes from one-day old adult males were dissected in PBS, incubated in Accutase (Stemcell Technologies, #07920) for10 minutes at room temperature to permeabilize the tissue, fixed in 4% formaldehyde/PBS with 0.1% Triton-X100 (PBST) for 10 minutes, incubated twice in 0.3% sodium deoxycholate/PBST for 10 minutes each^75^, incubated with primary antibodies in A+t buffer at 4^°^C overnight, and then with fluorescently-labeled secondary antibodies (1:200 dilution, Jackson ImmunoResearch). Testes were stained with 0.5 μg/mL DAPI/PBS and mounted in 80% glycerol on slides, and imaged on a Stellaris 8 confocal microscope (Leica) with 20X or 63X objectives. Pseudo-colored maximum-intensity projections were adjusted and composited in ImageJ, Adobe Photoshop, and Adobe Illustrator. For measuring signal intensities of the HLB in testes, we identified proliferative zone nuclei from gonial cell nuclei by bamGAL4-induced RFP fluorescence, and then used Photoshop to select the area of each HLB defined by MPM2 staining. We summed the RNAPII-S5p signal fluorescence pixel intensity of each HLB in unsaturated images, and normalized scores for each testis by the mean intensity of the proliferative zone, which is not affected by knockdowns in gonial cells.

### Imaging tissue culture cells

Drosophila *Kc167* and human K562 cells were swelled with a hypotonic 0.5% sodium citrate solution, then smashed onto glass slides in a Cytospin 4 centrifuge (Thermo). Slides were fixed with 4% formaldehyde/PBST and incubated with primary antisera in A+t buffer, then with fluorescently-labeled secondary antibodies (1:200 dilution, Jackson ImmunoResearch), stained with 0.5 μg/mL DAPI/PBS, mounted in 80% glycerol, and imaged by epifluorescence on an EVOS FL Auto 2 inverted microscope (Thermo Fisher Scientific) with a 40X objective. Pseudo-colored images were adjusted and composited in Adobe Photoshop and Adobe Illustrator.

### CUT&Tag chromatin profiling

To perform CUT&Tag^8^, we dissected 20 imaginal wing discs from male 3rd instar larvae in PBS buffer, and transferred them to a tube containing Accutase (Stemcell Technologies, #07920) at 25^°^C for 30 minutes. We then added an equal volume of 30% BSA to block proteases, and ran the material through a 30 ½ gauge needle once to dissociate tissue. Tissue suspensions were divided between 4-8 reaction tubes, and bound with BioMag Plus ConA (Bangs Inc., #531) magnetic beads. Tissue culture samples in media were added directly to ConA beads for binding, and then lightly fixed onto beads with 0.1% formaldehyde at room temperature for 1’. All samples were incubated with the following CUT&Tag solutions sequentially: primary antibodies diluted in Wash+ buffer (20 mM HEPES pH 7.5, 150 mM NaCl, 0.5 mM spermidine, 0.05% triton-X100, 2 mM EDTA, 1% BSA, with Roche cOmplete protease inhibitor) overnight at 4^°^, secondary antibodies (in Wash+ buffer) for 1 hour at room temperature, and then incubated with pAGTn5 (Epicypher 15-1017) in 300Wash+ buffer (20 mM HEPES pH 7.5, 300 mM NaCl, 0.5 mM spermidine, 0.05% triton-X100 with cOmplete protease inhibitor) for 1 hour. After one wash with 300Wash+ buffer, samples were incubated in 300Wash+ buffer supplemented with 10 mM MgCl_2_ for 1 hour at 25^°^C to tagment chromatin. Tissue culture samples were tagmented in CUTAC buffer (10 mM TAPS pH 8.5, 20% DMF) supplemented with 5 mM MgCl_2_ to enhance tagmentation efficiency. Samples were washed with 10 mM TAPS pH 8.5 and DNA was released with 0.1% SDS, 0.012 U/µL thermolabile protease K (NEB P8111S) in 10 mM TAPS at 37^°^ and inactivated at 55^°^C. Libraries were enriched with 14 cycles of PCR as described [Kaya-Okur et al 2019], and sequenced in dual-indexed paired-end mode (PE50) on the Illumina NextSeq 2000 or NovaSeq platforms at the Fred Hutchinson Cancer Center Genomics Shared Resource. Paired-end reads were mapped to this assembly using Bowtie2 using parameters, e.g.: --end-to-end --very-sensitive --no-mixed --no-discordant -q --phred33 -I 10 -X 700.

### Gene score tables

To summarize the enrichment of profiling features across histones, we counted mapped reads from the start to the end of each gene in.bam files using subreads/feature_counts with option ‘_o’. Counts for each replication-coupled histone were summed, and counts for all genes were scaled by total mapped reads to give Counts per Million (CPM) reads. These values are provided in **Supplementary Table 2**.

### Genomic display

Files of mapped reads were converted to genome coverage with bedtools/bamcoverage^76^ and displayed in the UCSC genome browser^77^. Selected regions were exported as PDFs and formatted with Adobe Illustrator.

## Data availability

Sequencing data have been deposited in GEO under accession code GSE280833. Data for NPAT profiling in K562 cells (SH_Hs_NPA1_20190217, SH_Hs_NPA2_20190217, SH_Hs_NPA4_20190217, SH_Hs_NPA8_20190217, SH_Hs_NPB1_20190217, SH_Hs_NPB2_20190217, SH_ Hs_NPB4_20190217, and SH_Hs_NPB8_20190217 was previously published^8^, and were merged here into one file (SH_Hs_NPB4_20190217). Data for RNAPII-S5p profiling in K562 cells (SH_Hs_K5xlin_PolS5P_3cy_0320, SH_Hs_ K5xlin_PolS5P_6cy, SH_Hs_K5xlin_PolS5P_9cy_0320, SH_Hs_K5xlin_PolS5P_12cy_0320, and SH_Hs_K5xlin_ PolS5P_20k_0320 were previously published^46^, and were merged here in one file (SH_Hs_K5xlin_PolS5P_20k_0320).

## Acknowledgements

We thank Amanda Amodeo and Yuki Shindo for *histone-Dendra2* reporter lines including two unpublished lines: the *His2ADendra2* HRU reporter and the *His2AVDendra2* reporter, Bob Duronio for histone locus rescue lines, Christine Codomo and Jorja Henikoff for technical and bioinformatic support, and Paul Talbert for pointing out the significance of viral histone H4 protein.

## Author contributions

K.A. and S.H. conceived the study. K.A., M.W., B.N.T., and V.V. performed the experiments. K.A., S.H., and V.V. performed data analysis. K.A. wrote the manuscript. K.A., S.H., and X.C. reviewed and edited the manuscript. All authors approved the manuscript.

## Competing interests

S.H. and K.A. have filed patent applications on related work.

## Supplementary Information

**Supplementary Table 1. Sample IDs and sequencing results**.

Attached spreadsheet.

**Supplementary Table 2. Histone gene count tables**.

Attached spreadsheet.

**Supplementary Table 3. Differential gene expression between wildtype and** Δ***HisC*; *12XWT* wing imaginal discs**.

Attached spreadsheet.

## Bioinformatics for figures

**For Figure 1:** mapped reads for Mute (BT2579), Mxc (BT3065), Rpb1 (BT542), RNAPII-S5p (BT1781), and RNAPII-S2p (BT3067) were converted to bigwigs with bedtools/genome coverage and displayed in the UCSC genome browser.

**For Figure 2: (a)**.bam files were counted using subReads/ featureCounts with option “-o” across a list of all annotated UCSC knownGenes in the dm6 genome assembly. Reads for each histone subtype were summed for each genotype, and counts in each histone subtype in the 12X line were divided by counts in the wildtype line. **(b)** Mapped reads for Mxc (BT3065), Mute (BT3064), H3K4me1 (BT3072), H3K4me2 (BT3068), H3K4me3 (BT3074), H3K27ac (BT3069), H3K36me3 (BT826), H3K9me1 (BT1777), H3K9me2 (BT1778), H3K9me3 (BT1175), H3K27me3 (BT1173), uH2A (BT827), and IgG (BT828) were converted to bigwigs with bedtools/genome coverage and displayed in the UCSC genome browser.

**For Figure 5i:** Mapped reads for Mapped reads for H3K-27ac (BT3505), RNAPII-S5p (BT3417), Mxc (BT3195), and histone H4 (BT3510) were converted to bigwigs with bedtools/genome coverage and displayed in the UCSC genome browser.

**For Figure 8: (b**,**c)** K562 tracks Mapped reads for H3K27ac (BT3514), RNAPII-S5p (SH K5xlin_PolS5P_0320), NPAT (SH NPAT_0217_pool), and histone H4 (BT3518) were converted to bigwigs with bedtools/genome coverage and displayed in the UCSC genome browser. **(d)** Gene scores were counted from.bam files for K562 cell profiling using subReads/featureCounts with option “-o” across a list of all annotated genes in the hg19 (February 2009) genome assembly from -200 bp of annotated gene starts to gene ends. Histone genes are displayed shaded from maximum to minimum counts for each epitope.

**Supplementary Figure 1.**
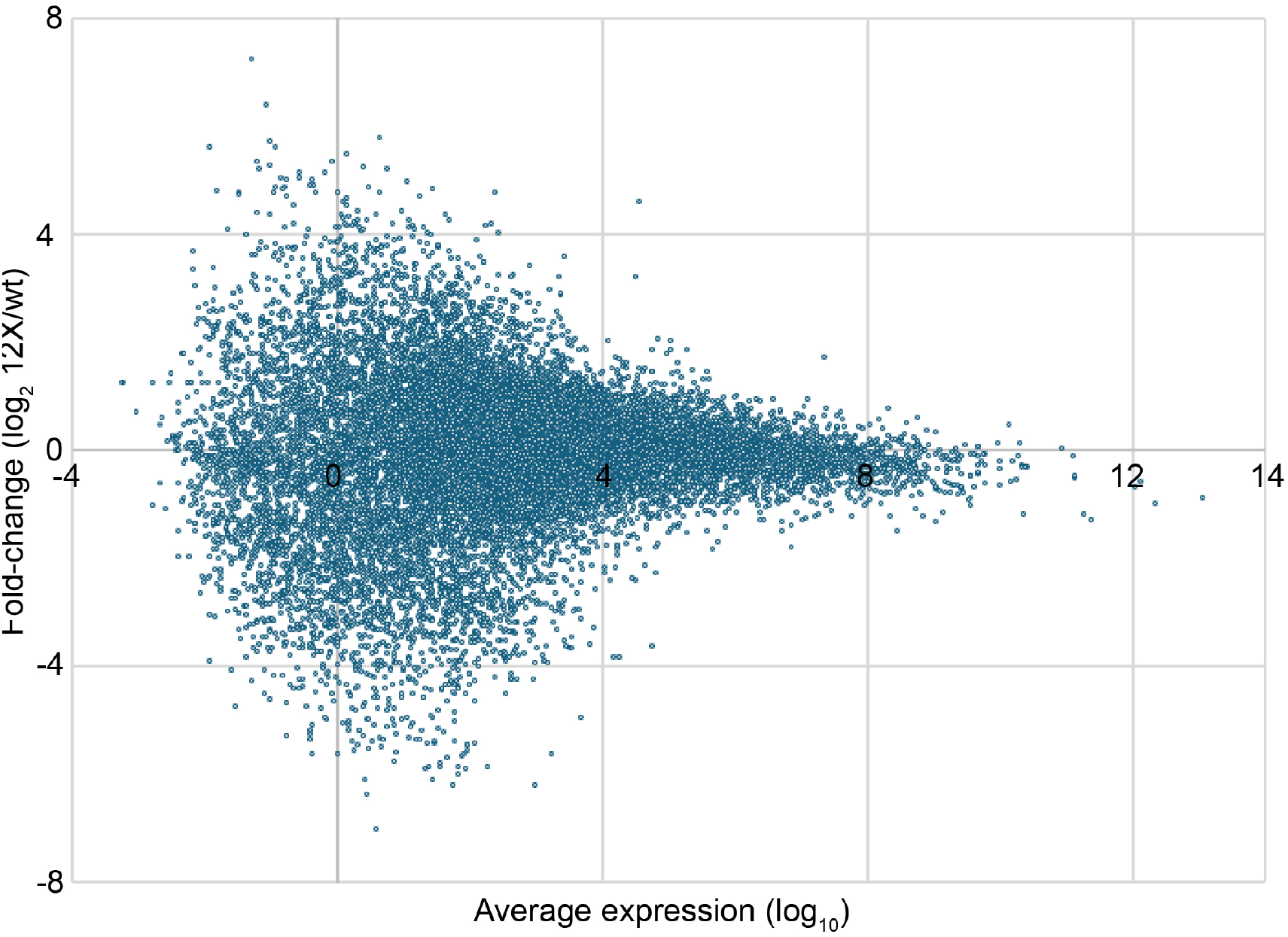
MA plot of RNAPII-S2p signal at genes in wildtype and *HisCΔ; 12XWT* wing imaginal discs. No genes with significantly-different expression between the two genotypes is detected. Values are presented in Supplementary Table 3.

**Supplementary Figure 2.**
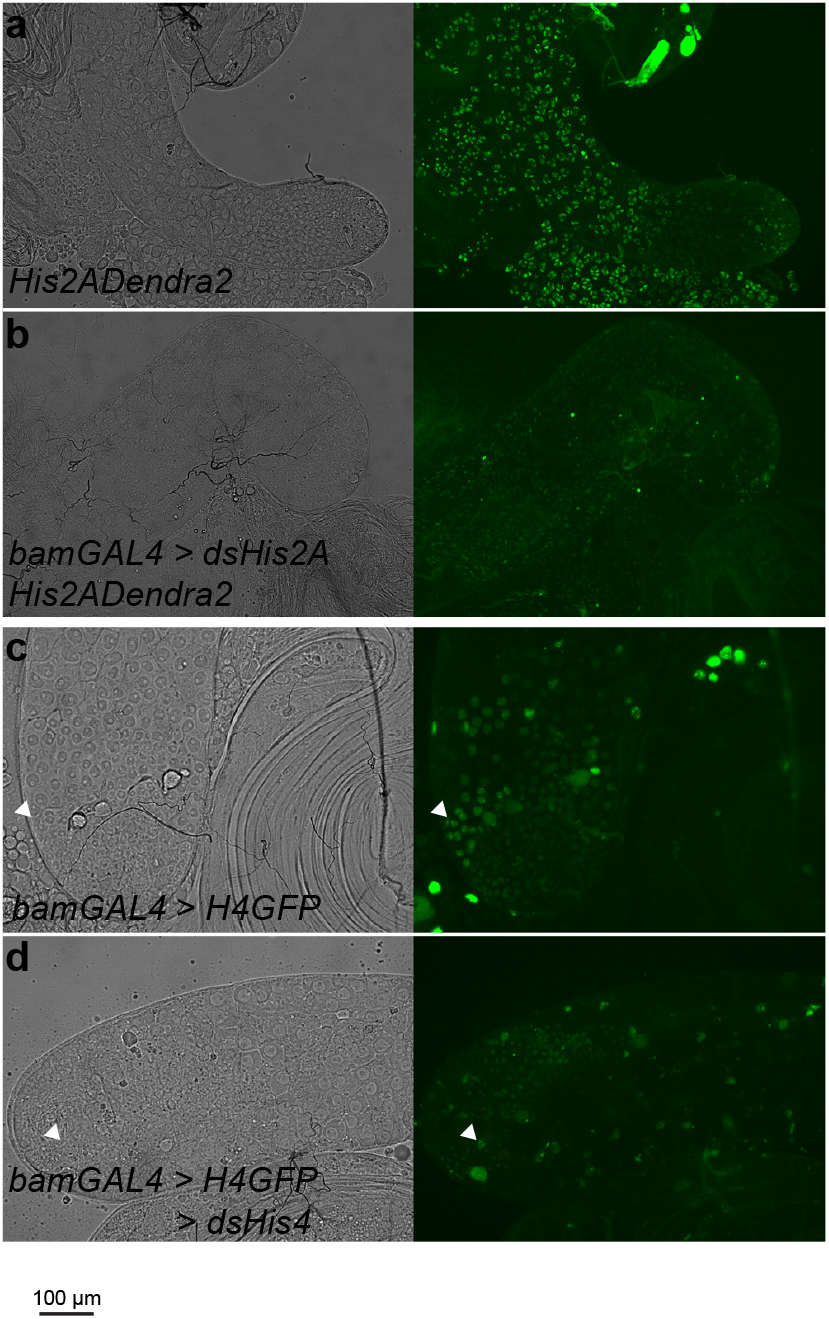
Knockdown efficiency of two histone RNAi constructs in the testis. **(a**,**b)** Identical exposures of testes from males carrying the *His2ADendra2* reporter. Normal males **(a)** show strong fluorescence in somatic cells of the apical tip and in developing primary spermatocytes. Males with *bamGAL4*-induced knockdown of *His2A* **(b)** have reduced fluorescence throughout the testis. **(c**,**d)** Identical exposures of testes from males carrying the *H4GFP* reporter. Males with *bamGAL4*-induced expression **(c)** show strong fluorescence a the HLB in gonial cells. Males with *bamGAL4*-induced induction of H4GFP and knockdown of *His4* **(d)** have reduced fluorescence with little or no HLB staining.

## Key Resources Table

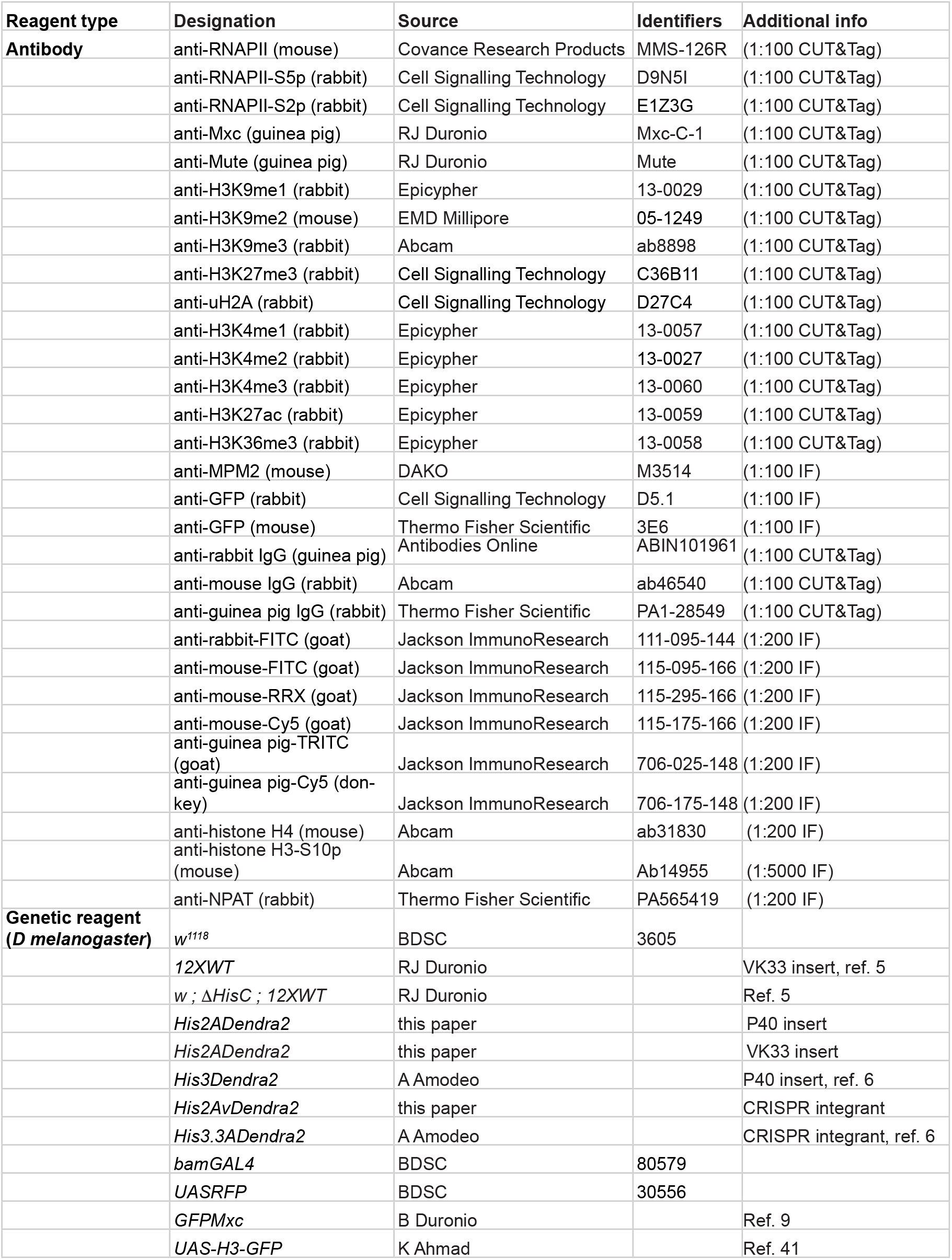
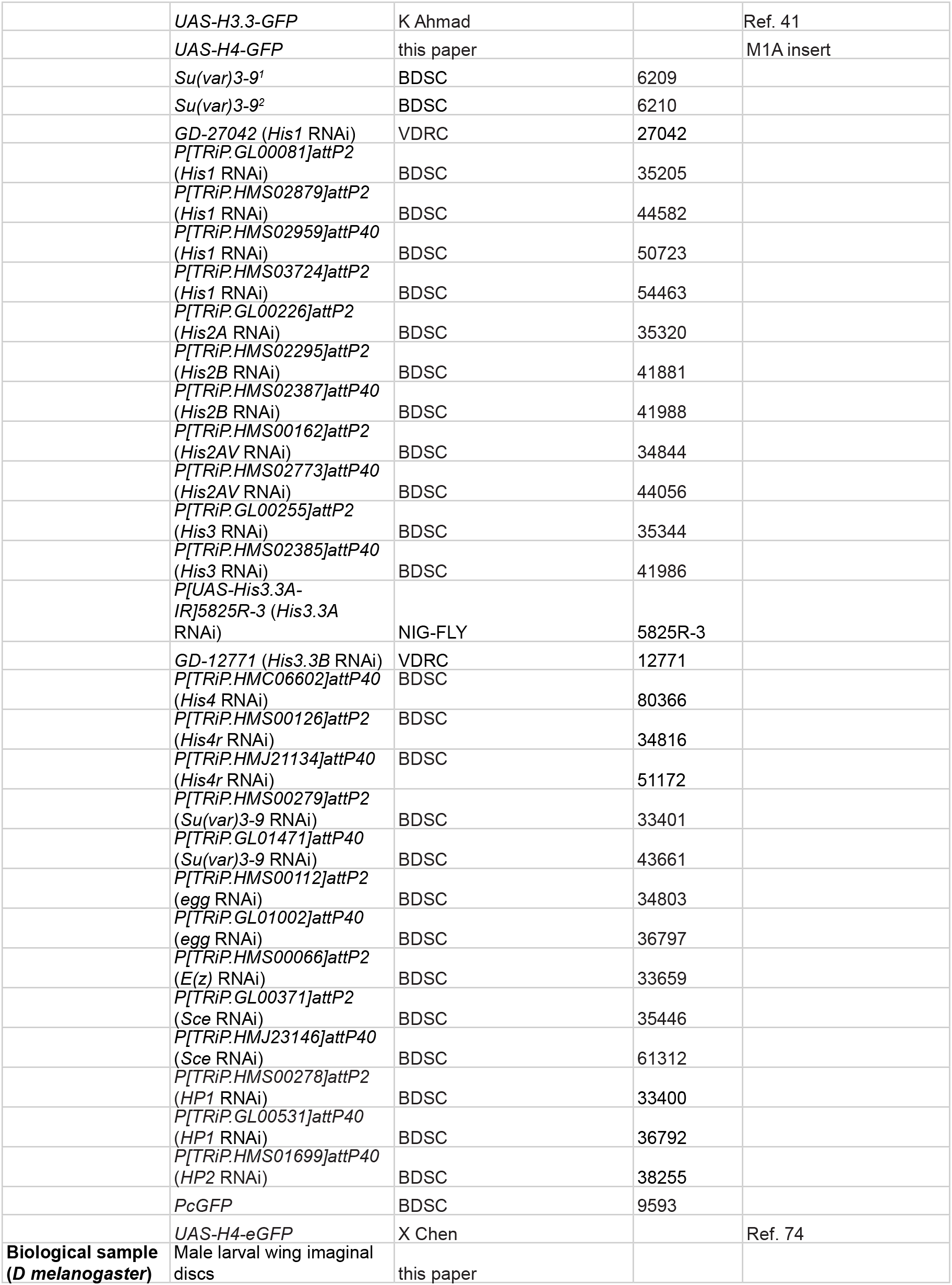
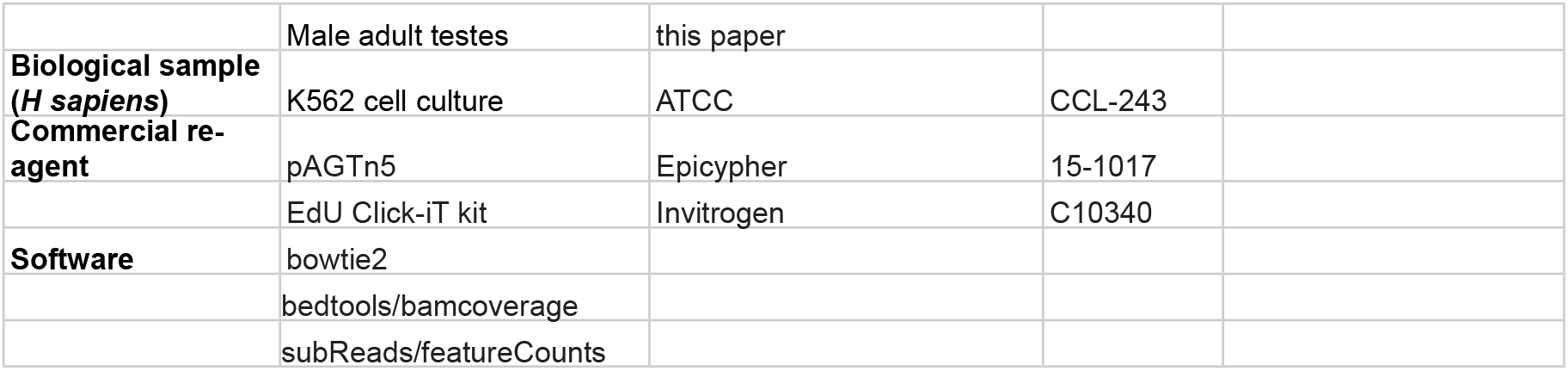

## Notes

### Summary of Updates

Additional analysis in Figure 4 and Figure 7, and a model proposed in Figure 8

### Funding

- Howard Hughes Medical Institute: S.H.
- NIH NIGMS: GM152821

