## Supplementary figures and images for "Cell-cycle-dependent repression of histone gene transcription by histone H4"

### Supplementary Figure 1

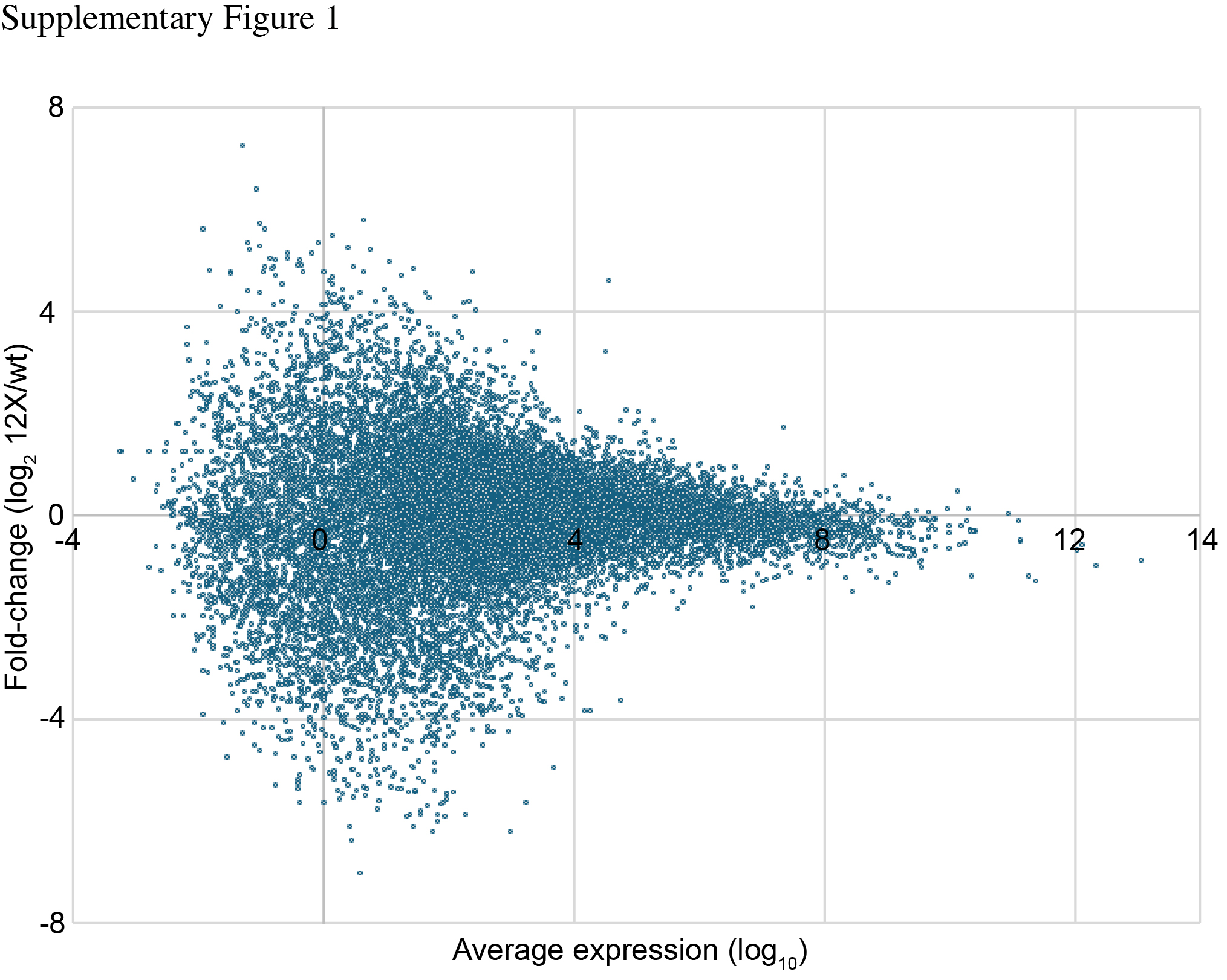

### Supplementary Figure 2

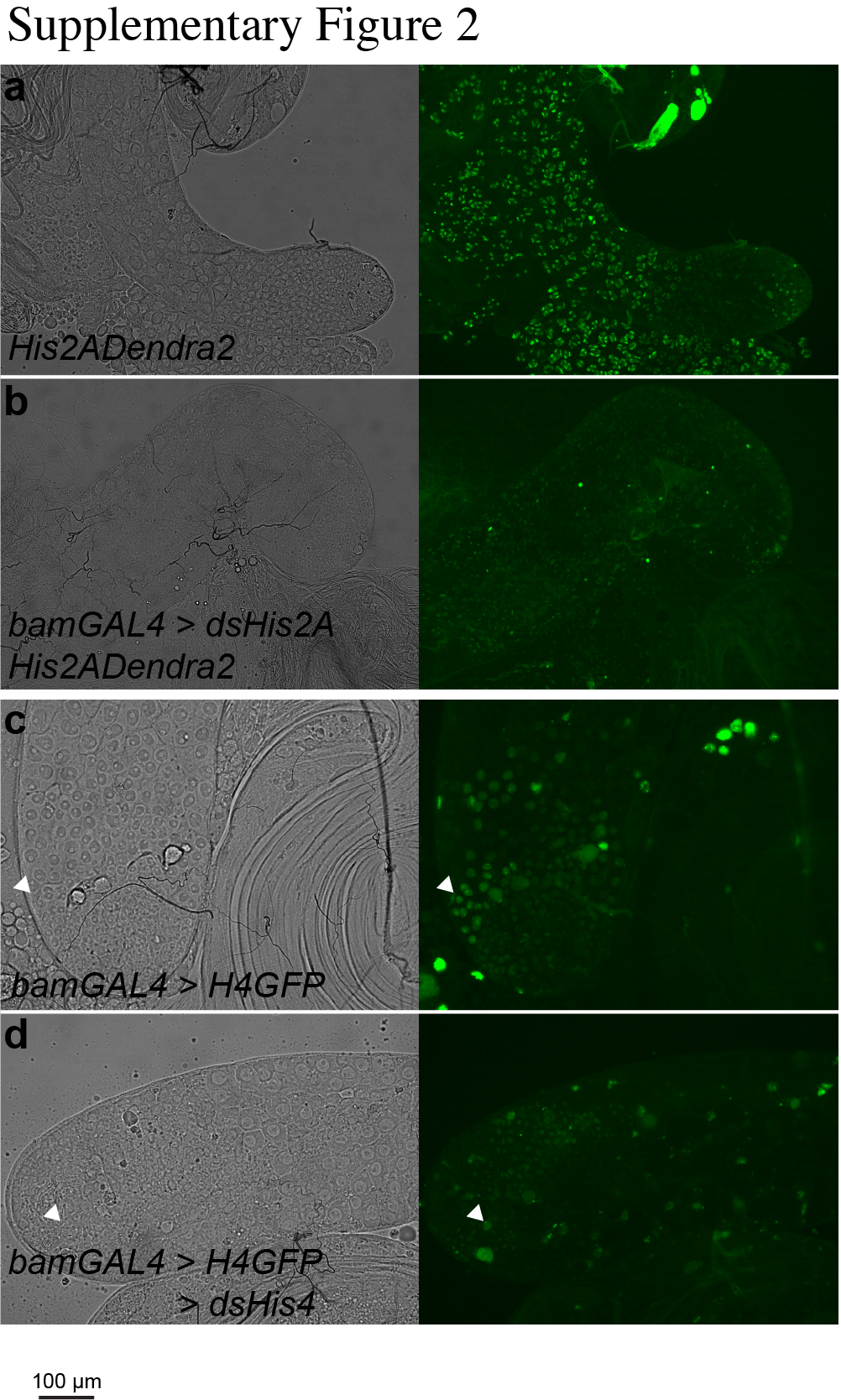
